# Description and biology of two new egg parasitoid species, *Trichogramma chagres* and *T. soberania* (Hymenoptera: Trichogrammatidae) reared from eggs of Heliconiini butterflies (Lepidoptera: Nymphalidae: Heliconiinae) collected in Panama

**DOI:** 10.1101/493643

**Authors:** Jozef B. Woelke, Viktor N. Fursov, Alex V. Gumovsky, Marjolein de Rijk, Catalina Estrada, Patrick Verbaarschot, Martinus E. Huigens, Nina E. Fatouros

## Abstract

Two new minute egg parasitoid wasp species belonging to the genus *Trichogramma* (Hymenoptera: Trichogrammatidae), *T. chagres* sp. nov. and *T. soberania* sp. nov., were found in a tropical lowland rainforest in Panama, Central America. In this paper, we describe, illustrate and discuss the biology, morphological and molecular characterization of the two new *Trichogramma* wasp species. Both species were collected from eggs of passion vine butterflies, *Agraulis vanillae vanillae* (Lepidoptera: Nymphalidae: Heliconiinae) and unidentified Heliconiini species, laid on different *Passiflora* species (Malpighiales: Passifloraceae). A female *T. soberania* sp. nov. wasp was noted on the wings of a female *Heliconius hecale melicerta* butterfly caught in the wild. This suggests that this species may occasionally hitch a ride on adult female butterflies to find suitable host eggs. Our study adds two more species identifications to the scarce record of *Trichogramma* wasps from the widespread Heliconiini butterflies in Central America.

## Introduction

The Passion-vine or longwing butterflies belonging to the tribe Heliconiini (Lepidoptera: Nymphalidae: Heliconiinae), which comprise the genus *Heliconius* Kulk and related genera, are a highly diverse group of butterflies from the Neotropics (Brown 1981; Gilbert 1991; Harvey 1991; Penz and Peggie 2003). Heliconiini butterflies are important study objects to unravel the coevolution between insects and (host) plants (Brown 1981; Ehrlich and Raven 1964). Passion vine or *Passiflora* (Malpighiales: Passifloraceae) plants are exclusively used as host plants for their offspring (Benson et al. 1975; Brown 1981). Eggs are typically brightly coloured and laid singly or in (small) groups on new shoots, tendrils or older leaves (Benson et al. 1975; Brown 1981).

Many nymphalid butterflies are toxic and, especially those belonging to the genus *Heliconius*, are well known model organisms for studies on Müllerian mimicry, with numerous species converging to a common wing pattern (Brown 1981; Gilbert 1991; Jiggins et al. 2004; Mavárez et al. 2006). *Heliconius* butterflies feed on pollen, which allows them to live up to six months (Gilbert 1972). Despite the well-known ecological interactions and evolutionary history of Heliconiini butterflies, only very few studies have identified some of their natural enemies to species level (Guerrieri et al. 2010; Querino and Zucchi 2002, 2003a,b; Zhang et al. 2005; Zucchi et al. 2010). We conducted a study of the egg parasitoid community of Heliconiini butterflies on *Passiflora* plants in a tropical lowland rainforest in Panama and found that most species of parasitic wasps were not yet described. Currently, only the newly found species *Ooencyrtus marcelloi* (Hymenoptera: Encyrtidae) has been described from this study (Guerrieri et al. 2010).

*Trichogramma* wasps (Hymenoptera: Trichogrammatidae) are minute (± 0.5 mm long) gregarious egg parasitoids that are used worldwide as important biological control agents against agricultural pest insects, predominantly Lepidopterans (Li 1994; Nagarkatti and Nagaraja 1977; Polaszek 2010; Smiths 1996; Stinner 1977). In addition to Lepidoptera, *Trichogramma* can occasionally also parasitize eggs of Coleoptera, Diptera, Hemiptera, Neuroptera, Megaloptera and other Hymenoptera (Nagarkatti and Nagaraja 1977; Polaszek 2010).

*Trichogramma* species are considered as polyphagous parasitoids, although the level of polyphagy may differ between species (Romeis et al. 2005; Zucchi et al. 2010). They mainly use chemical information to find suitable host eggs (Fatouros et al. 2008). For example, some wasps spy on the anti-aphrodisiac pheromone emitted by mated female Pierid butterflies and hitch a ride on the latter to oviposition sites. Such a chemical-espionage-and-ride strategy has been shown to be used also by other egg parasitoid species (Fatouros et al. 2005; Huigens et al. 2010; Fatouros and Huigens 2012; Huigens and Fatouros 2013). Anti-aphrodisiac pheromones were also identified in 11 *Heliconius* species (Estrada et al. 2011). We hypothesize that *Trichogramma* wasps can also use anti-aphrodisiacs to find *Heliconius* butterflies to hitch-hike and parasitize their freshly laid eggs.

Worldwide, around 210 *Trichogramma* species have been described (Pinto 2006). The diversity of species is well-described for North America where 60 species of *Trichogramma* are recorded (Pinto 1999, Zucchi et al. 2010). In South America 41 species of *Trichogramma* are described of which most are recorded in Brazil (28 species) followed by Venezuela (13), Colombia (9) and Peru (7) (Parra and Zucchi 2004; Velasquez de Rios and Teran 1995, 2003; Querino et al. 2017; Querino and Zucchi 2003a,b, 2005; Zucchi 1988; Zucchi et al. 2010). In Central America 22 species of *Trichogramma* are recorded and among them only one species, *T. panamense* Pinto has been described from Panama (Pinto 1999), showing that our two new species are the second and third species records for this country.

Morphological species identification of the complex *Trichogramma* genus can be done by observing male genitalia, which are a very useful diagnostic character (Nagarkatti and Nagaraja 1971). Females are often impossible to identify to species level by morphological characteristics. Stouthamer et al. (1999) developed a molecular tool for *Trichogramma* species identification based on the Internal Transcribed Spacer ITS-2 region of ribosomal DNA that is now-a-days, together with morphology, often used for species identification. Here, we describe two new species of *Trichogramma* on the basis of species morphology with illustrations and the sequence of the ITS-2 DNA, as well as the biology of the two species. They were reared from Heliconiini eggs that were deposited on several species of *Passiflora* plants. Moreover, we monitored Heliconinii butterflies for the presence of hitch-hiking *Trichogramma* wasps.

## Materials and methods

### Wasp collection

*Trichogramma* wasps were collected from Heliconiini eggs (Figure 1a,b) during a field survey from February until April 2008 in a tropical lowland rainforest in Soberania National Park (Parque Nacional Soberanía) (Figure 1c), and in the town of Gamboa and surroundings, in the Republic of Panama. In our research area, six *Passiflora* plant species were found: *Passiflora foetida* L. *var. isthmia* Killip (Figure 1d), *P. auriculata* Kunth, *P. vitifolia* Kunth (Figure 1e), *P. biflora* Lamarck (Figure 1f), *P. coriacea* Juss., and *P. menispermifolia* Kunth. Plants were labelled and checked once or twice a week for the presence of Heliconiini eggs. Eggs were put separately into small glass vials and taken to a laboratory of the Smithsonian Tropical Research Institute (STRI) in Gamboa. Eggs were checked daily for the presence of caterpillar or wasp emergence. If a caterpillar emerged, it was reared or released onto the location where the egg was collected or into STRI butterfly facilities. Parasitized eggs become black several days before wasp(s) emergence and can then be differentiated from unparasitized eggs. *Trichogramma* wasps were sexed upon emergence and then offered *Heliconius erato* or *H. melpomene* and later *Mamestra brassicae* (Lepidoptera: Noctuidae) eggs to rear more individuals for identification and study the wasp’s biology. For detailed biological studies, fresh and developing eggs of *M. brassicae* were offered to the females of each strain. Behavior of oviposition was observed. Eggs were dissected on certain time intervals, so details of the immature development and morphology were revealed using different stereo- and upright microscopes.

**Figure 1.**
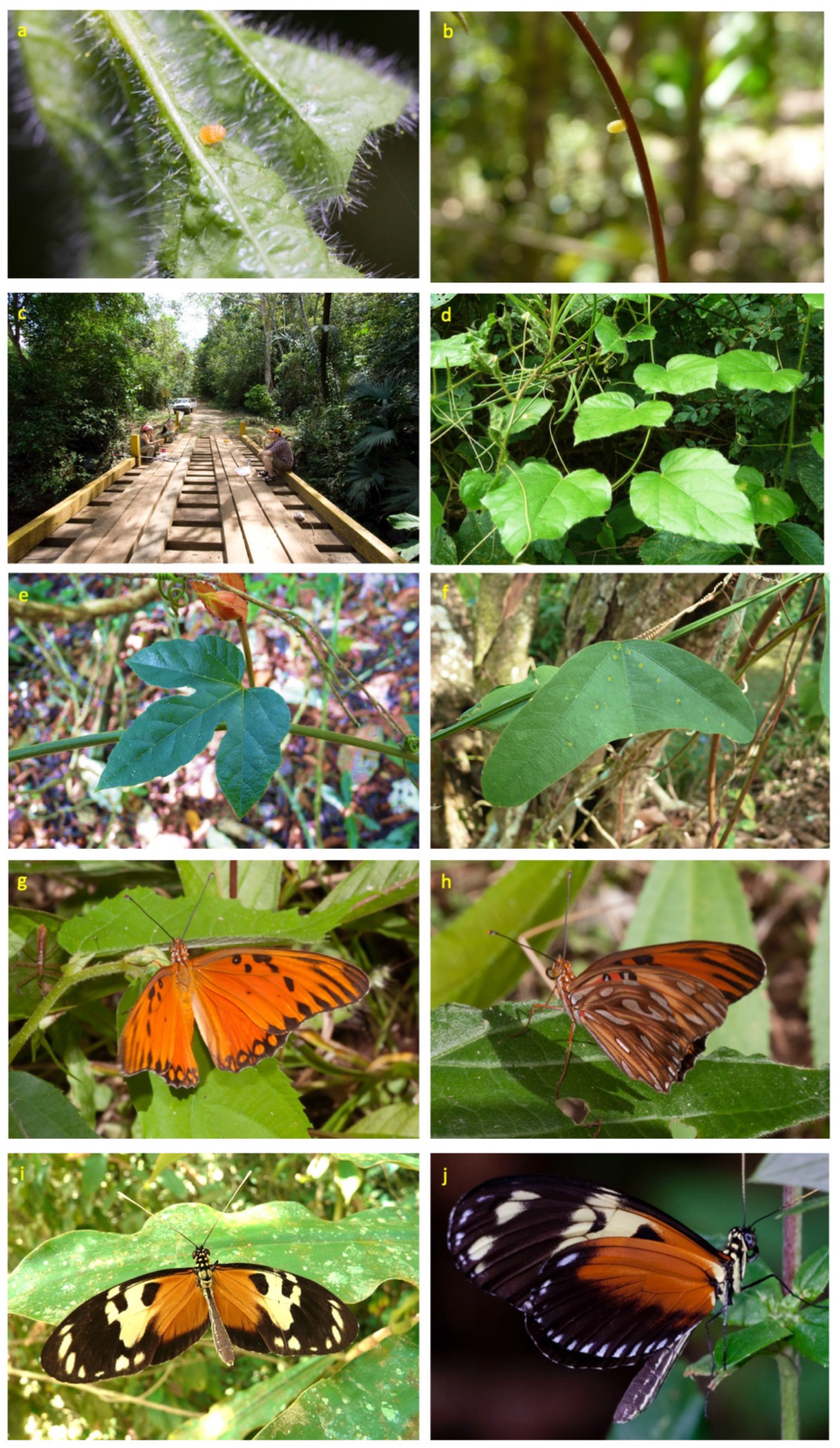
(a) Heliconiini egg on a new shoot of *Passiflora foetida var. isthmia;* (b) Heliconiini egg on tendril of *P. vitifolia;* (c) field site, pipeline road in Soberania National Park; (d) *Passiflora foetida var. isthmia* on which both new *Trichogramma* species were found from Heliconiini eggs; (e) *Passiflora vitifolia* on which *T. chagres* sp. nov. was found from a Heliconiini egg; (f) *Passiflora biflora* on which *T. soberania* sp. nov. was found from a Heliconiini egg; (g-h) *Agraulis vanillae vanillae*, host of both new *Trichogramma* species; (i-j) *Heliconius hecale melicerta*, on which a *T. soberania* sp. nov. wasp was found.

Besides the collections of eggs, we also caught Heliconiini adults to check for the presence of hitch-hiking wasps on their bodies. The Heliconiini species known in the research areas are *Agraulis vanillae vanillae* (Figure 1g-h), *Dione juno huascuma, Dryas iulia moderata, Eueides aliphera gracilis, E. lybia olympia; Heliconius cydno chioneus, H. doris viridis, H. erato petiverana, H. hecale melicerta* (Figure 1i-j), *H. hecalesia, H. ismenius, H. melpomene rosina, H. sapho sapho, H. sara fulgidus* and *Philaethria dido* (Estrada and Jiggins 2002; Naisbit 2001; Dr. Annette Aielo personal information). Butterflies were collected with a butterfly net. Inside the net, a plastic pot attached with a rubber band was placed to avoid wasp escape (as described in Fatouros and Huigens 2012). When a butterfly was caught, it was cooled down to decrease its activity. After collection, all butterflies were placed overnight in a refrigerator (4°C) at the STRI laboratory in Gamboa and checked the next day carefully for the presence of parasitic wasps. Butterflies were fed with honey water and released in the same area where they were caught.

Wasps were stored in 95% ethanol and shipped to the Schmalhausen Institute of Zoology of the National Academy of Sciences, Ukraine, for morphological identification or to the Laboratory of Entomology, Wageningen University and Research, The Netherlands for molecular identification.

### Morphological species identification

The descriptions, measurements and figures were made with the aid of an Olympus CX-40 microscope. Photographs were taken using the software Olympus Capture. The terminology and abbreviations of morphological structures and ratios of male genitalia and antenna are follows Pinto (1999) and Burks and Heraty (2002). Descriptions are based on holotype and paratype male specimens, which were mounted on glass slides. The measurements of body parts were based on five male specimens for each of both species described below. Body parts were measured in micrometers (μm) and summarized in the description as range followed by the mean in parenthesis (avg means average).

Some of the commonly used morphological terms were abbreviated as follows: length of aedeagus (AL, Figure 2a), length of basal part of aedeagus (AL-B, Figure 2a), length of dorsal aperture (DA-L), length (DLA-L, Figure 2a) and width (DLA-W, Figure 2a) of dorsal lamina (DLA), maximum width (GW, Figure 2a) or length (GL, Figure 2a) of genital capsule (GC), width of basal part of genital capsule (GW-B, Figure 2a), apical distance (AD, Figure 2a), length (IVP-L, Figure 2a) and width (IVP-W, Figure 2a) of intervorsellar process (IVP), internal length of parameres (PL, Figure 2a), length of the longest seta of flagellum (SL), and maximum width of flagellum (FW), marginal vein of fore wing (MV).

We calculated the ratios of (1) SL/FW; (2) GL/GW, (3) GW/GW-B, (4) DA-L/GL, (5) DLA-L/DLA-W, (6) GW/DLA-W, (7) DLA-L/GL, (8) IVP-L/IVP-W, (9) AD/IVP-L, (10) AD/GL, (11) PL/DLA-L, (12) AL/GL, and (13) AL-B/AL.

### Molecular species identification based on ITS-2 gene

Wasp species identification was also performed based on the ribosomal ITS-2 gene as described previously (Fatouros and Huigens 2012; Gonçalves et al. 2006; Huigens et al. 2004; Stouthamer et al. 1999).

#### DNA extraction

From every vial, with emerged wasp(s) of one Heliconiini egg, one wasp was taken and dried on a filter paper. Every wasp was crushed separately in a 0.5 ml Eppendorf tube with a closed Pasteur pipette. After crushing, 50 μl of Chelex solution (5%) and 4 μl of proteinase K (20mg/ml) were added. Finally, the samples were incubated overnight at 56°C followed by 10 min at 95°C.

#### PCR amplification

A PCR reaction was performed for every sample. In a 0.2 ml Eppendorf tube 25 μl volume consisting of 18.43 μl of distilled water, 2.5 μl 10x PCR reaction buffer (HT Biotechnologies Ltd., Cambridge, UK), 2.5 μl DNA template, 0.5 μl dNTP (10mM), 0.5 μl forward primer (25μM), 0.5 μl reverse primer (25μM) and 0.07 μl Taq polymerase (5 units/μl) (HT Biotechnologies Ltd., Cambridge, UK) was added. To amplify the ITS-2 region the forward primer 5’-TGTGAACTGCAGGACACATG-3’ and the reverse primer 5’-GTCTTGCCTGCTCTGAG-3’ were used. The PCR cycling program was 3 min at 94°C, 33 cycles of 40 s at 94°C, 45 s at 53°C and 45 s at 72°C, followed by 10 min at 72°C after the last cycle. PCR products were run on a 1.5% agarose gel and stained with ethidium bromide.

#### Cloning and sequencing

Each amplified ITS-2 gene was cloned and sequenced. ITS-2 products were purified from the gel using the MinElute Gel Extraction Kit (QIAGEN GmbH, Hilden, Germany) for DNA fragment purification. The PCR fragment was ligated to a pGEM-T vector (Promega, Madison, WI, USA) and transformed into *Escherichia coli* XL2blue cells (Stratagene, La Jolla, CA, USA). Correct insertion of the ITS-2 fragments was confirmed by PCR. To purify the plasmid, the GenElute Plasmid Miniprep Kit (Sigma-Aldrich Chemie GmbH, Steinheim, Germany) was used. By using an Applied Biosystems automatic sequencer the ITS-2 fragments were sequenced. ITS-2 sequences were finally aligned and matched against sequences present in GenBank and those present in the large database of Prof. Dr. R. Stouthamer (University of California, Riverside, USA).

### Species deposition

The holotype and series of paratypes specimens of both species (mounted on glass slides) are deposited in the collection of Schmalhausen Institute of Zoology of National Academy of Sciences of Ukraine, Kiev, Ukraine (SIZK). Paratypes are deposited in the collection of Natural History Museum, London, UK (BMNH) and Naturalis Biodiversity Center, Leiden, The Netherlands (RMNH).

## Results

In total, we collected 317 singly laid eggs of different Heliconiini species. From these 317 eggs, we found 3.2% being parasitized by *Trichogramma* wasps and 1.6% per *Trichogramma* species described here. From those parasitized eggs, 33 females and 7 males of the first species (later named *T. chagres*), and 27 females and 6 males of the second species (later named *T. soberania*) emerged. We also collected eight egg clusters, with a total of 996 eggs, of the gregarious species *Dione juno*. None of the *D. juno* eggs were parasitized. Moreover, we caught 133 butterflies of the Heliconiini tribe on which we found one *Trichogramma* wasp in the net (later named *T. soberania*), which was presumably hitchhiking on the butterfly body. After studying the collected materials, we conclude that both *Trichogramma* species are new to science. The descriptions are given below.

## Taxonomy

> Family **Trichogrammatidae** Haliday, 1851
>
> Genus **Trichogramma** Westwood, 1833
>
> ***Trichogramma chagres*** Fursov and Woelke, sp. nov.

**Figure 2.**
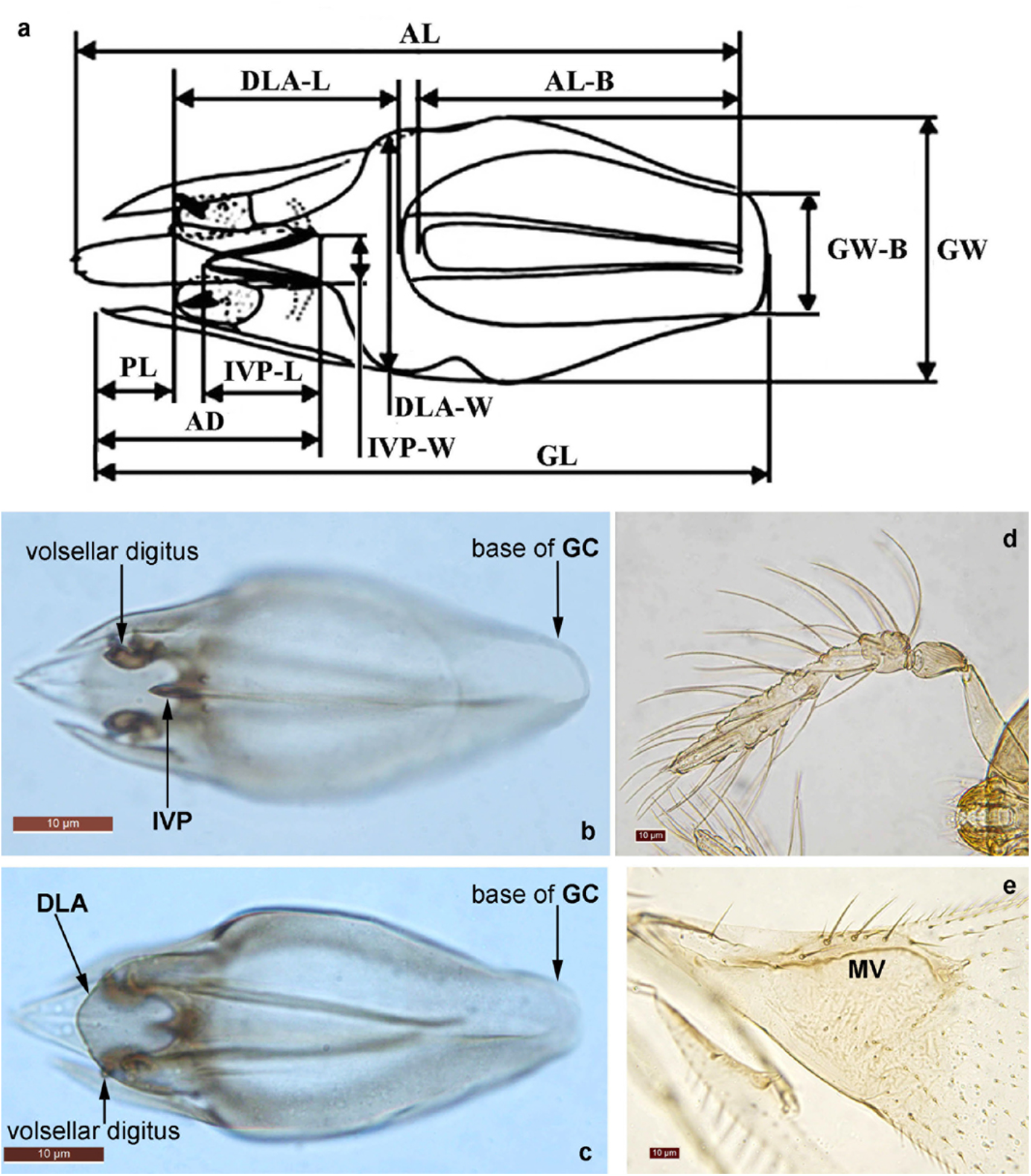
*Trichogramma chagres* sp. nov., holotype male: (a) structures of male genitalia with abbreviations of measurements (explanation in text); (b) genitalia, ventral view; (c) genitalia, dorsal view; (d) antenna; (e) veins of fore wing.

**Figure 3.**
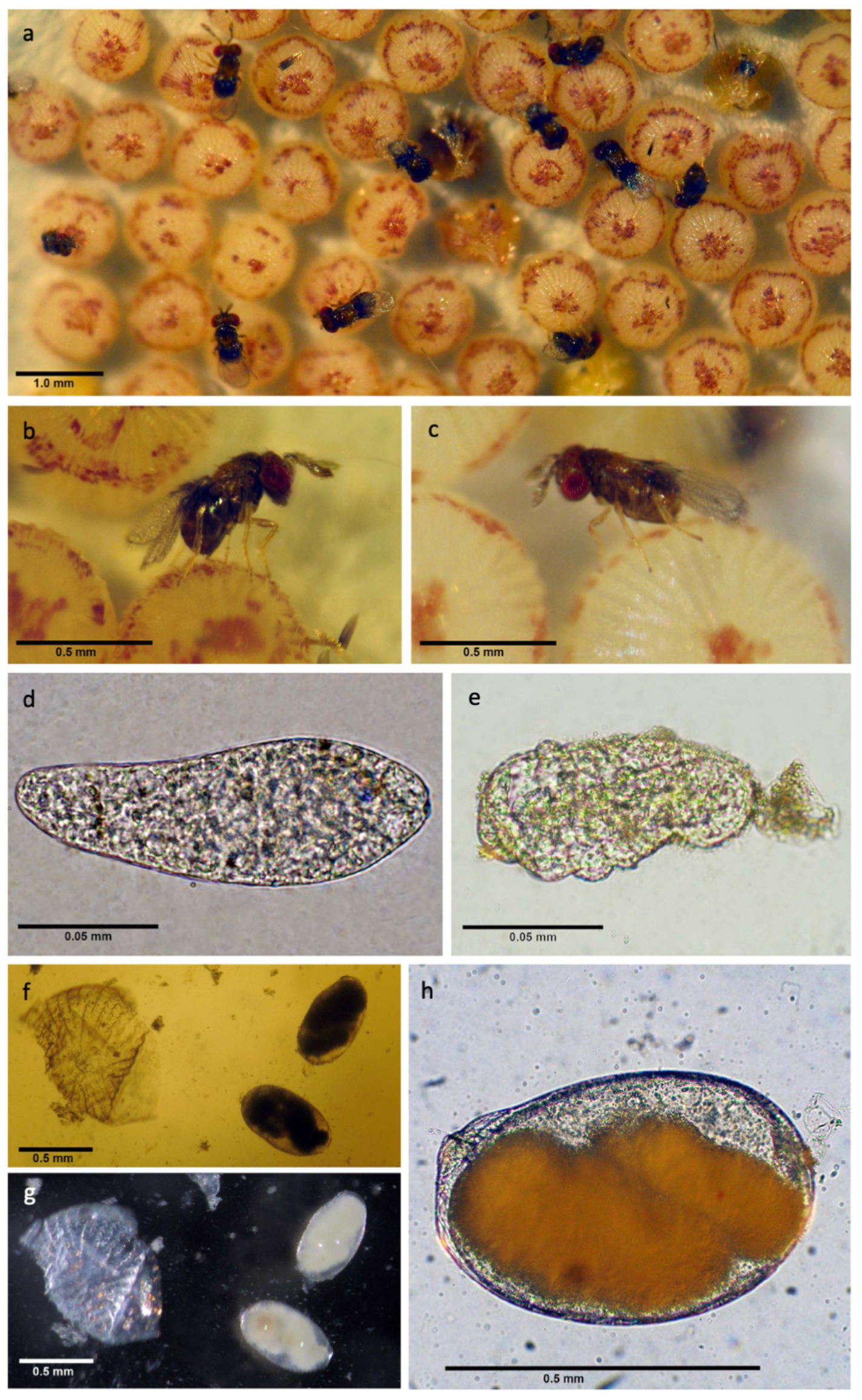
Biology of *Trichogramma chagres* sp. nov. strain L21: (a-c) female adult(s) parasitizing egg(s) of *Mamestra brassicae;* (d) freshly laid wasp egg; (e) newly hatched larva; (f-h) mature larva: (f-g) two mature larvae and chorion of consumed host egg in direct (f) and reflected (g) light, (h) habitus of mature larva with pulsing mid gut full of host egg yolk.

### Diagnosis

*Trichogramma chagres* sp. nov. is characterized by a wide GC (about 2.21–2.30 times as long as wide, Figure 2b-c), very wide DLA (Figure 2c), very long, narrow and apically sharp IVP (Figure 2b), long and sharp setae of antennae (about 1.92–2.11 times as long as width of clava, Figure 2c). The new species is morphologically close to *T. benetti* Nagaraja and Nagarkatti, *T. drepanoforum* Pinto and Oatman and *T. atopovirillia* Oatman and Platner, but it is well distinguishable from them all in the possession of the distinctly long IVP (about 1.21–1.57 times as long as wide, Figure 2c), which is much shorter in the other species. Apart from the shape of IVP, *T. chagres* sp. nov. differs from *T. bennetti* in having a very narrow base of GC (Figure 2b-c) (it is widened basally in *T. bennetti*). The new species is distinguishable from both *T. drepanoforum* and *T. benetti* in having the wide DLA being shaped as a spade with a subtriangular tip (Figure 2c) (the tip of DLA is evenly acute in both, *T. drepanoforum* and *T. benetti)*. Also, the tip of DLA is extended beyond the tips of vorsellar digiti in *T. chagres* sp. nov. (Figure 2b-c), unlike in *T. benetti*.

### Description

Based on holotype and 4 paratype male specimens.

Color of head and antennae yellow; meso- and metasoma dark brown, except bright yellow axillae, propodeum and base of gaster. All legs yellow, except hind femora and tibiae which are dirty yellow-brownish.

Antenna (Figure 2d) with flagellum 5.26–6.51 (avg. 5.67) times as long as maximum width, 1.92–2.11 (avg. 1.95) times as long as length of scape; SL/FW = 2.85–3.43 (avg. 3.10). Number of flagellar setae 35–38 (Figure 2d).

GL (Figure 2b-c) 112.25–137.18 (avg. 122.73), GW 50.87–62.11 (avg. 55.50), DA-L 82.15–105.89 (avg. 90.03). DLA-L 23.62–38.29 (avg. 30.85), DLA-W 29.39–33.14 (avg. 31.66); IVP-L 6.97–13.22 (avg. 8.76), IVP-W 4.62–9.90 (avg. 6.18); AD 25.51–38.86 (avg. 31.83); PL 25.81–38.29 (avg. 30.80); AL-B 42.13–44.94 (avg. 44.69); AL 85.63–116.85 (avg. 97.39). GC wide, with wide DLA, GL/GW = 2.21–2.30 (avg. 2.24), but very narrow at the base (Figure 2b-c), widest medially or subapically (at distance of 0.53 of GL), then sharply narrowed to the top, with elongated dorsal aperture. DA-L/GL = 0.70–0.78 (avg. 0.74). DLA very wide, spade-shaped, without basal lobes, but with small sharp lateral-apical notches, with nearly parallel lateral sides and with rounded and slightly sharpened apical part (Figure 2c), extended over apical parts of parameres (Figure 2b-c). DLA-L/DLA-W = 0.81–1.18 (avg. 0.98). GW/DLA-W = 1.60–1.92 (avg. 1.77). DLA-L/GL = 0.47–0.50 (avg. 0.49). Apex of DLA not extending beyond apical part of parameres, but extending beyond apices of vorsellar digiti (Figure 2b). IVP sclerotized, large, with wide base and with sharp teeth-like apex (Figure 2b). IVP-L/IVP-W = 1.21–1.57 (avg. 1.42). AD/IVP-L = 2.58–5.19 (avg. 3.76). AD/GL = 0.23–0.31 (avg. 0.27). Apical part of GC narrowed gradually, without curvature. Parameres extending to the apex of vorsellar digiti at a distance 1.56–2.75 (avg. 2.07) as long as IVP (Figure 2b). PL/DLA-L = 0.77–1.28 (avg. 1.04). DA-L/GL = 0.71–0.78 (avg. 0.75). AL/GL = 0.47–0.50 (avg. 0.49). AL-B/AL = 0.73–0.86 (avg. 0.82). Wings. Fore wings transparent. MV with three large and four small setae (Figure 2e).

### Material examined

**Holotype male** (SIZK), Panama, Pipeline Road, 9°08’31.8”N, 79°43’30.6”W, collected 11^th^ March 2008 from egg of Heliconiini butterfly (Lepidoptera: Nymphalidae: Heliconiinae) found on *Passiflora foetida var. isthmia* (Malpighiales: Passifloraceae) (coll. J.B. Woelke and M. de Rijk), specimen on glass slide under 2 small cover slips (genitalia under right side cover slip), covered by black pen, on slide No 2019 (strain L21) (in Canada balsam).

**Paratypes** same label (all from strain L21), 1 female on slide No 2019 (SIZK); 1 male and 1 female on slide No 2020 (SIZK); 1 male and 1 female on slide No 2021 (RMNH); 1 male and 1 female on slide No 2022 (BMNH) (all in Canada balsam).

**Additional material** (SIZK) same label (strain L23, this parasitized Heliconiini egg was collected on the same plant, date and location as strain L21), 5 males and 4 females on slide No 1875; 3 males and 3 females on slide No 1876; 3 males and 3 females on slide No 1877; 3 males and 3 females on slide No 1878 (all in Canada balsam).

Panama, Gamboa, 9°07’05.8”N, 79°41’41.1”W, collected 3 April 2008 from egg of Heliconiini butterfly (Lepidoptera: Nymphalidae: Heliconiinae) found on *P. vitifolia* (Malpighiales: Passifloraceae) (coll. J.B. Woelke and M. de Rijk), strain L31 (SIZK), 3 males and 2 females on slide No 1871; 3 males and 3 females on slide No 1872; 3 males and 3 females on slide No 1873; 5 males and 4 females on slide No 1874 (all in Canada balsam).

**Field records** Panama, Pipeline Road, 9°08’31.8”N, 79°43’30.6”W, #112, 7 females and 2 males collected 26 February 2008 from egg of *Agraulis vanillae vanillae* (Lepidoptera: Nymphalidae: Heliconiinae) found on *Passiflora foetida var. isthmia* (Malpighiales: Passifloraceae) (coll. J.B. Woelke and M. de Rijk);

Panama, Pipeline Road, 9°08’31.8”N, 79°43’30.6”W, #295 (origin of strain L21) 10 females and 2 males collected 11 March 2008 from egg of Heliconiini butterfly (Lepidoptera: Nymphalidae: Heliconiinae) found on *Passiflora foetida var. isthmia* (Malpighiales: Passifloraceae) (coll. J.B. Woelke and M. de Rijk);

Panama, Pipeline Road, 9°08’31.8”N, 79°43’30.6”W, #300, 6 females and 1 male collected 11 March 2008 from egg of Heliconiini butterfly (Lepidoptera: Nymphalidae: Heliconiinae) found on *Passiflora foetida var. isthmia* (Malpighiales: Passifloraceae) (coll. J.B. Woelke and M. de Rijk);

Panama, Pipeline Road, 9°08’31.8”N, 79°43’30.6”W, #301 (origin of strain L23) 10 females and 2 males collected 11 March 2008 from egg of Heliconiini butterfly (Lepidoptera: Nymphalidae: Heliconiinae) found on *Passiflora foetida var. isthmia* (Malpighiales: Passifloraceae) (coll. J.B. Woelke and M. de Rijk);

Panama, Gamboa, 9°07’05.8”N, 79°41’41.1”W, #1039 (origin of strain L31), unknown number of wasps collected 3 April 2008 from egg of Heliconiini butterfly (Lepidoptera: Nymphalidae: Heliconiinae) found on *P. vitifolia* (Malpighiales: Passifloraceae) (coll. J.B. Woelke and M. de Rijk).

### Host

Wasps were reared from eggs of *Agraulis vanillae vanillae* (Figure 1g-h) and Heliconiini spp. found on *Passiflora foetida* L. *var. isthmia* Killip (Figure 1d) and *P. vitifolia* Kunth (Figure 1e).

### Biology

Idiobiont endoparasitoid. All specimens of this species were reared from collected eggs of Heliconiini butterflies, which were deposited on *Passiflora* plants. The collected wasps had an average of 8.25 ± 2.06 SD females and 1.75 ± 0.50 SD males per egg and having a sex-ratio of 21.21%.

More specific information about strain L21 (Figure 3), L23 and L31 (Figure S1). Females actively oviposit into fresh and relatively mature host eggs (with red bands) (Figure 3a-c, S1a-b). The freshly laid parasitoid egg is about 0.08-0.09 mm long (Figure 3d, S1c), developing embryo within the egg (24 h after oviposition) is about 0.15 mm long and 0.05 mm wide (in its widest part) (Figure S1d-e). The newly hatched larva is about 0.1 mm long, with distinct head capsule bearing mandibles and three thoracic segments separated by deep constrictions, unsegmented abdomen and a caudal formation behind (Figure 3e). The mature fed-up larva is about 0.7 mm long, 0.4 mm wide, with pulsing mid gut full of consumed host yolk, with remnants of caudal bladder (membranes) behind (Figure 3f-h, S1g-h). No molts were traced during the larval development, and the mandibles of hatching and mature larvae are of the same size, about 0.01 mm long (Figure S1f).

### Distribution

Panama, tropical lowland rainforest of the Soberania National Park (Parque Nacional Soberanía) (Figure 1c), and in the town of Gamboa and surroundings.

### Etymology

The Chagres (in Spanish: *Rio Chagres*) is the largest river (193 km) in the Panama Canal’s watershed. The river that is surrounding our research areas, making a sharp bend around the town of Gamboa.

### Sequence analysis

MegaBLAST analysis revealed that our Internal Transcribed Spacer 2 (ITS-2) sequences of *T. chagres* sp. nov. matched with 40% query cover and 89% identity to *T. chilotraeae* Nagaraja and Nagarkatti in GenBank. Sequence ID *T. chagres* sp. nov: MK159692.

> ***Trichogramma soberania*** Fursov and Woelke, sp. nov.

**Figure 4.**
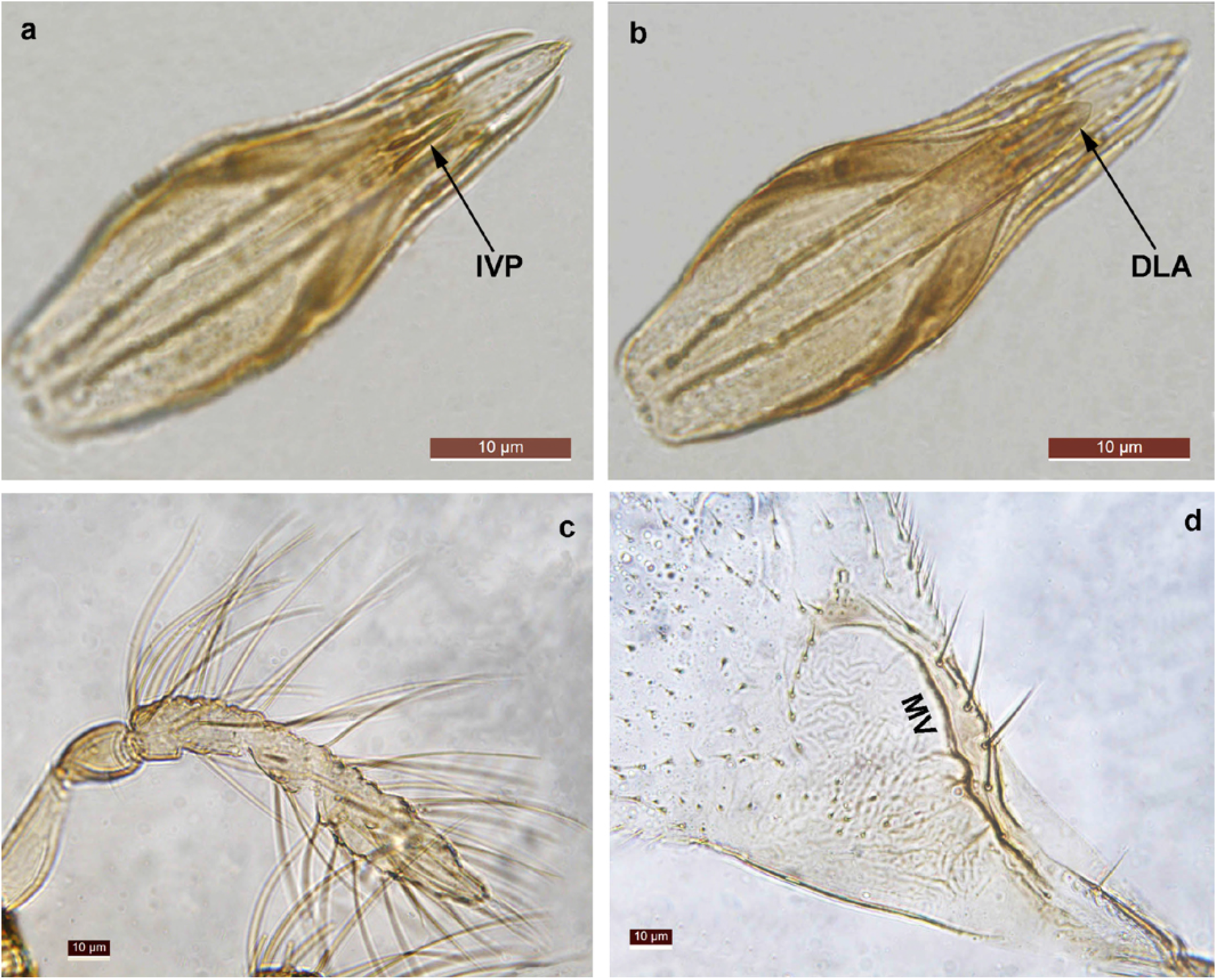
*Trichogramma soberania* sp. nov., holotype male (a) genitalia, ventral view; (b) genital, dorsal view; (c) antenna; (d) veins of fore wing.

**Figure 5.**
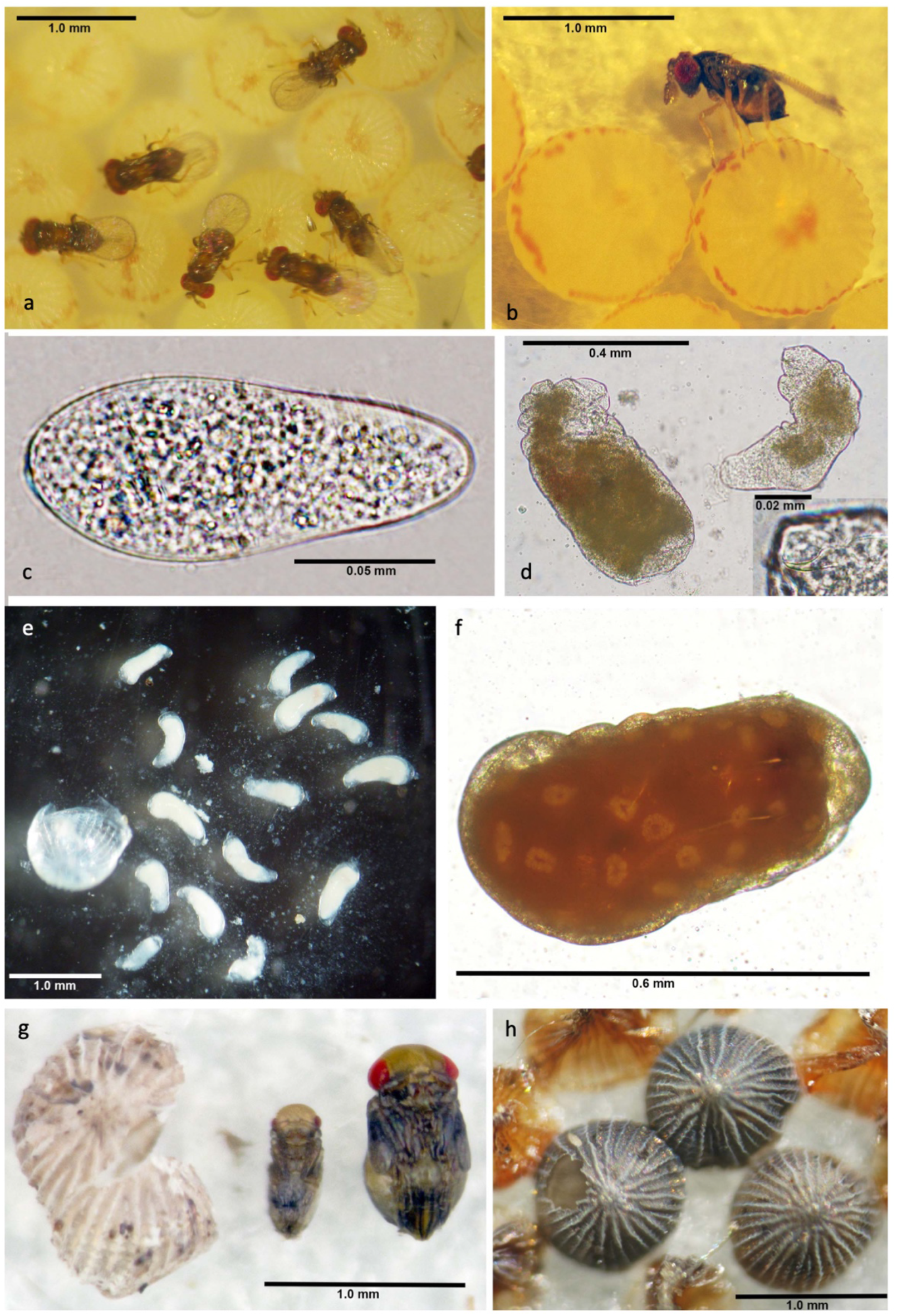
Biology of *Trichogramma soberania* sp. nov. strain L20 (a-b) female adult(s) parasitizing egg(s) of *Mamestra brassicae;* (c) freshly laid wasp egg; (d) 48 h larva; (e) 14 larvae can develop inside a *M. brassicae* egg; (f) fully-grown larva; (g) male and female pupae; (h) *Mamestra brassicae* eggs turn black after consumption or parasitism.

### Diagnosis

*Trichogramma soberania* sp. nov. is characterized by a narrow shape of phallobase (about 2.5 times as long as wide, Figure 4a-b), very narrow and apically sharp and elongated DLA (Figure 4b), long and sharp setae of antennae (about 2.5 times as long as width of clava (Figure 4c). The species is morphologically close to *T. exiquum* Pinto and Platner and *T. pretiosum* Riley but can be differentiated from both species by the presence of a long and very narrow IVP (2.0–3.62 times as long as wide, Figure 4a). Also, *T. soberania* sp. nov. is discernible from *T. pretiosum* in the shape of DLA (Figure 4b), which is notably narrower in the latter species.

### Description

Based on holotype and 8 paratype male specimens.

Color of head, antennae, meso- and metasoma dark brown, except light yellow scutellum, propodeum and base of metasoma, all legs brown, eyes red.

Antenna (Figure 4c) with flagellum 5.40–6.19 (avg. 5.83) times as long as its maximum width, and 1.95–2.08 times (avg. 2.02) as long as length of scape; SL/FW = 2.71–3.04 (avg. 3.0). Number of flagellar setae 37–44 (Figure 4a).

GL 112.60–158.55 (avg. 132.05), GW 35.33–61.02 (avg. 50.96); GW-B 20.46–24.58 (avg. 22.54); DA-L 62.10–95.54 (avg. 76.47).

DLA 36.11–46.03 (avg. 43.50), width 30.08–48.35 (avg. 38.84) (Figure 4b); IVP-L 9.51–18.00 (avg. 14.70); AD 31.20–42.41 (avg. 35.60); PL 29.87–35.08 (avg. 32.82); AL-B 70.58–88.83 (avg. 75.73); AL 126.35–167.28 (avg. 144.22).

GC very narrow basally, widest medially or subapically, and then again sharply narrowed apically; with elongate dorsal aperture. DA-L/GL = 0.56–0.61 (avg. 0.59). GL/GW = 2.52–3.31 (avg. 2.80). GW/GW-B 1.73–2.49 (avg. 2.26). DLA sharply narrowed medially and smoothly narrowed apically, subtriangular, with distinct basal lobes, small sharp lateral notches, and with smoothly rounded apical part (Figure 4b). Basal lobes of DLA not extended to lateral sides of GC. Apex of DLA not extended beyond apical part of parameres, but extended beyond the apex of volsellae and volsellar digiti, as well as beyond the apex of IVP (Figure 4a-b). Apex of DLA narrower than width of aedeagus (Figure 4b). DLA-L/DLA-W = 0.29–0.40 (avg. 0.34). GW/DLA-W = 1.24–1.75 (avg. 1.41). DLA-L/GL 0.40-0.56 (avg. 0.34). IVP sclerotized on both lateral sides, long, with wide base and with very narrow awl-like sharpened apex (Figure 4a). IVP not extended beyond the apex of vorsellar digiti, and beyond the apex of DLA. IVP-L/IVP-W = 2.0–3.62 (avg. 2.57). AD/IVP-L = 2.03–3.76 (avg. 2.42). AD/GL = 0.27–0.42 (avg. 0.31). Apical part of GC narrowed gradually, without curvature (Figure 4a-b). PL/DLA-L 0.63–0.73 (avg. 0.72). AL/GL = 1.04–1.09 (avg. 1.07). AL-B/AL = 0.48–0.56 (avg. 0.53).

Wings. Fore wings transparent, MV with four large and four small setae (Figure 4d).

### Material examined

**Holotype male** (SIZK), Panama, Pipeline Road, 9°08’31.8”N, 79°43’30.6”W, collected 4^th^ March 2008 from egg of Heliconiini butterfly (Lepidoptera: Nymphalidae: Heliconiinae) found on *Passiflora foetida var. isthmia* (Malpighiales: Passifloraceae) (coll. J.B. Woelke and M. de Rijk), specimen on glass slide under 3 small cover slip (genitalia under right left side cover slip), covered by black pen, on slide No 2023 (strain L20) (in Canada balsam).

**Paratypes** same label (all from strain L20), 1 male on slide No 2024 (SIZK); 1 male and 1 female on slide No 2025 (SIZK); 1 male and 1 female on slide No 2026 (SIZK); 1 male and 1 female on slide No 2027 (RMNH); 1 male and 1 female on slide No 2031 (BMNH); 1 male on slide No 2028 (SIZK); 1 male on slide No 2029 (SIZK); 1 male on slide No 2030 (SIZK) (all in Canada balsam).

**Additional material** (SIZK) same label (strain L20), 3 males and 1 female on slide No 1867; 3 males and 3 females on slide No 1868; 3 males and 2 females on slide No 1869; 3 males and 3 females on slide No 1870 (all in Canada balsam).

**Field records** Panama, Pipeline Road, 9°08’31.8”N, 79°43’30.6”W, #105, 7 females and 1 male collected 26 February 2008 from egg of *Agraulis vanillae vanillae* (Lepidoptera: Nymphalidae: Heliconiinae) found on *Passiflora foetida var. isthmia* (Malpighiales: Passifloraceae) (coll. J.B. Woelke and M. de Rijk);

Panama, Pipeline Road, 9°08’31.8”N, 79°43’30.6”W, #168, 10 females and 1 male collected 4 March 2008 from egg of Heliconiini butterfly (Lepidoptera: Nymphalidae: Heliconiinae) found on *Passiflora foetida var. isthmia* (Malpighiales: Passifloraceae) (coll. J.B. Woelke and M. de Rijk);

Panama, Pipeline Road, 9°08’31.8”N, 79°43’30.6”W, #180 (origin of strain L20), 2 females and 1 male collected 4 March 2008 from egg of Heliconiini butterfly (Lepidoptera: Nymphalidae: Heliconiinae) found on *Passiflora foetida var. isthmia* (Malpighiales: Passifloraceae) (coll. J.B. Woelke and M. de Rijk);

Panama, Plantation Road, 9°04’32.1”N, 79°39’32.3”W, #283, 5 females and 2 males collected 6 March 2008 from egg of Heliconiini butterfly (Lepidoptera: Nymphalidae: Heliconiinae) found on *Passiflora biflora* (Malpighiales: Passifloraceae) (coll. J.B. Woelke and M. de Rijk);

Panama, Pipeline Road, 9°08’31.8”N, 79°43’30.6”W, #346, 2 females and 1 male collected 20 March 2008 from egg of *Agraulis vanillae vanillae* (Lepidoptera: Nymphalidae: Heliconiinae) found on *Passiflora foetida var. isthmia* (Malpighiales: Passifloraceae) (coll. J.B. Woelke and M. de Rijk).

**Hitch-hiking record** Panama, Pipeline Road, 9°08’31.8”N, 79°43’30.6”W, #32, 1 female wasp was collected on a female *Heliconius hecale melicerta* butterfly, 12 February 2008 (coll. J.B. Woelke and M. de Rijk).

### Host

Wasps were reared from eggs of *Agraulis vanillae vanillae* (Figure g-h) and Heliconiini spp. found on *Passiflora biflora* Lamarck (Figure 1f) and *P. foetida* L. *var. isthmia* Killip (Figure 1d).

### Biology

Idiobiont endoparasitoid. All specimens of this species were reared of collected eggs of Heliconiini butterflies, which were deposited on *Passiflora* plants. The collected wasps had an average of 5.20 ± 3.42 SD females and 1.20 ± 0.45 SD males per egg and having a sex-ratio of 23.08%. Our finding of one female wasp on an adult female *Heliconius hecale melicerta* butterfly (Figure 1i-j) suggests that this species may occasionally hitch a ride on adult female butterflies to find suitable host eggs.

More specific information about strain L20. Female wasps oviposit several eggs into a *M. brassicae* host of various ages (fresh and mature with red stripes) (Figure 5a-b). Both winged and brachypterous females oviposit. Freshly laid egg is ovoid, about 0.12 mm long, and ~0.06 mm wide (Figure 5c). About 24 h after, an embryo is traceable within the egg. The freshly hatched larva is about as long as the egg, with poorly sclerotized mandibles. 48 h later the larva is already measured 0.4-0.6 mm long, has distinct mid gut full of consumed host yolk (Figure 5d). About 50 h after oviposition the entire egg is consumed, and larvae reach nearly their final size. Up to 14 larvae can develop in same egg of *M. brassicae* (Figure 5e) The fully-grown larva is about 0.7-0.8 mm long, swollen, with full mid gut; occasionally it is reddish with white spots (Figure 5f). No molt or changes in mandible size were traced during the larval development. The male and female pupae differ in size and colour pattern (Figure 5g). Host eggs turn black after consumption, likewise in case of parasitism by most *Trichogramma* species (Figure 5h).

### Distribution

Panama, tropical lowland rainforest of the Soberania National Park (Parque Nacional Soberanía) (Figure 1c), and in the town of Gamboa and surroundings.

### Etymology

Research was conducted in Soberania National Park (in Spanish: *Parque Nacion Soberanía*) in Panama, as well in the town of Gamboa, which is located in this park. The park is protected since 1980 and covers 220 km^2^ of tropical lowland rainforest.

### Sequence analysis

MegaBLAST analysis revealed that our Internal Transcribed Spacer 2 (ITS-2) sequences of *T. soberania* sp. nov. matched with 37% query cover and 93% identity to *T. chilotraeae* in GenBank. Sequence ID *T. soberania* sp. nov: MK159692.

## Acknowledgements

From February until April 2008 all wasp material was collected using permit no. SE/AP-3-08 and exported using permit nos. SEX/A-19-08 and SEX/A-32-08. All permits were provided by the Autoridad Nacional del Ambiente (ANAM) of Panama. We are grateful to Annette Aiello and Chris Jiggins for supporting the field survey, and Yde Jongema, John Pinto and Richard Stouthamer for their help in the early stages of wasp identifications. AG would like to thank Martijn Bezemer and Jeff Harvey for their support and advice provided throughout his stay at NIOO-KNAW and Wageningen University & Research.

---

**Figure S1.**
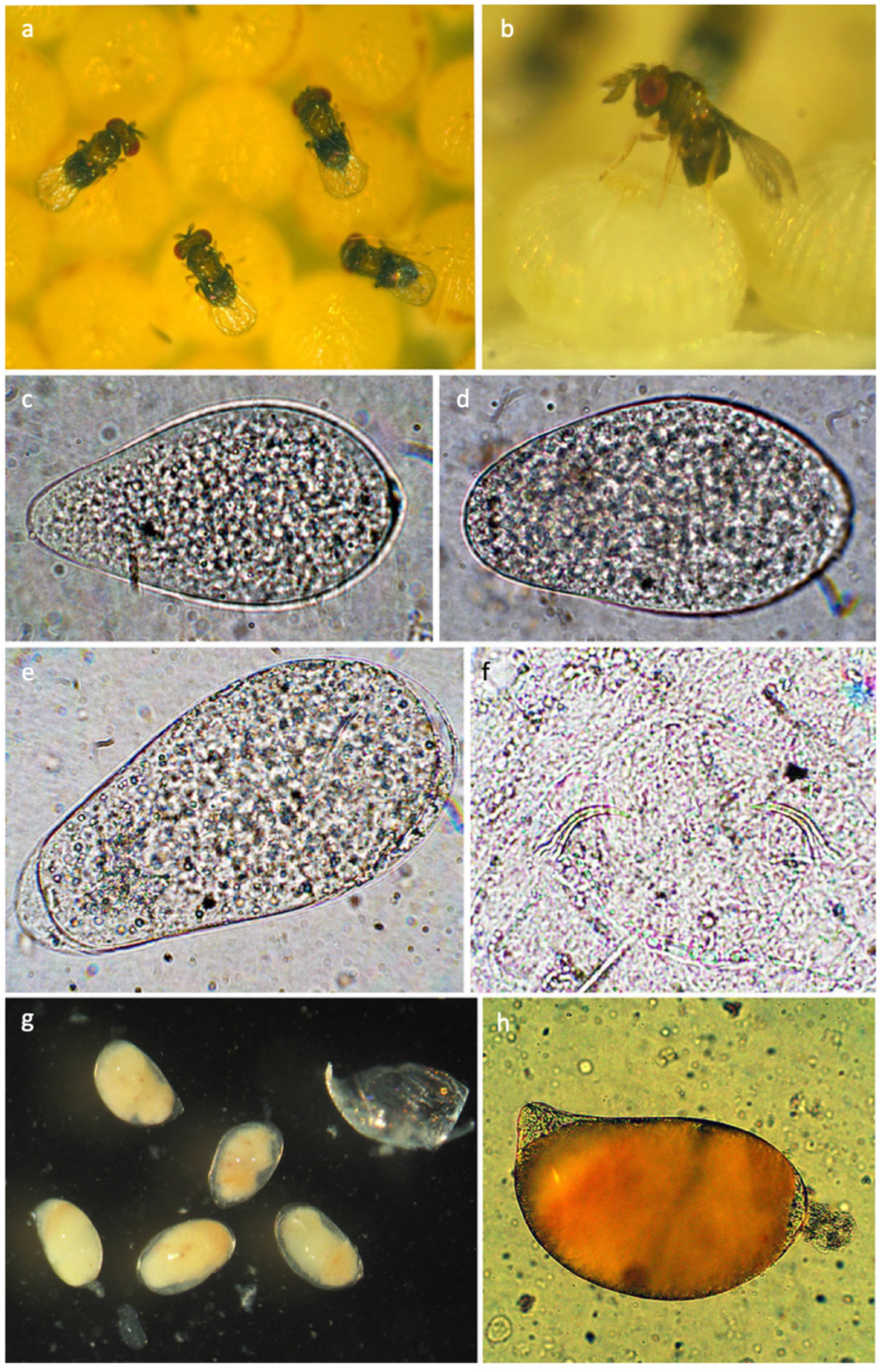
Biology of *Trichogramma chagres* sp. nov. strain L31 (a-b) female adult(s) parasitizing egg(s) of *Mamestra brassicae;* (c-e) eggs: (c) freshly laid egg; (d) developing egg; (e) egg with embryo inside; (f) mandibles of larva; (g) mature larvae isolated from consumed host egg and the egg’s chorion; (h) habitus of mature larva with pulsing mid gut full of host egg yolk.

